# CTRL: a label-free method for dynamic measurement of single-cell volume

**DOI:** 10.1101/817189

**Authors:** Kai Yao, Nash D. Rochman, Sean X. Sun

## Abstract

Measuring the physical size of the cell is valuable in understanding cell growth control. Current single-cell volume measurement methods for mammalian cells are labor-intensive, inflexible, and can cause cell damage. We introduce CTRL: Cell Topography Reconstruction Learner, a label-free technique incorporating Deep Learning and Fluorescence Exclusion for reconstructing cell topography and estimating mammalian cell volume from DIC microscopy images alone. The method achieves quantitative accuracy, requires minimal sample preparation, and applies to extensive biological and experimental conditions. Using this method, we observe a noticeable reduction in cell size fluctuations during cell cycle, which is consistent with the presence of a cell size checkpoint. (https://GitHub.com/sxslabjhu/CTRL)

## Main text

Cell size plays a critical role during cell growth, division, and proliferation^1–5^. Abnormalities in cell size regulation and growth control are thought to promote disease development^2,5–9^. Accurately measuring single-cell size remains a challenge for mammalian cells due to their irregular shape. Existing techniques require specialized hardware, fluorescent labeling^10,11^ and/or cell suspension^12–16^. Fluorescent labeling or over-expression of a target marker can alter cell function. Cell suspension alters the cell shape and biochemical signaling from the extracellular matrix, and also potentially affecting cell size. None of these methods has been successfully applied to measure mammalian cell growth at the single-cell level. While sensitive and accurate methods have been developed to measure single-cell mass over time^17^, the relationship between cell size and mass is not always clear.

An accurate and high throughput method of cell volume quantification is the Fluorescence Exclusion method (FXm), first proposed in 1983^14^ and subsequently developed and refined by several groups^18–20^. Cells are seeded in a micro-fabricated chamber and a membrane-impermeable high molecular weight fluorescent dye (e.g. FITC-dextran) is injected into the microchamber (Fig. 1a). The cell excludes its volume in the microchamber, therefore the total fluorescence loss is proportional to the cell volume. The FXm method obtains the cell volume from a single epi-fluorescence image, and therefore is high throughput^3,18–20^. However, due to endocytosis^3,20^ that is common in many cell types, the dye eventually enters the cytoplasm, and therefore FXm generally cannot accurately report cell volume in time-lapse without careful controls. Fluorescent imaging also introduces photobleaching, which alters the signal during time-lapse measurements. Moreover, microfluidic fabrication is needed to perform the experiment and the confinement of the microchamber may alter cell physiological processes over long periods. These drawbacks limit the use and applicability of FXm for studying cell growth.

**Fig. 1.**
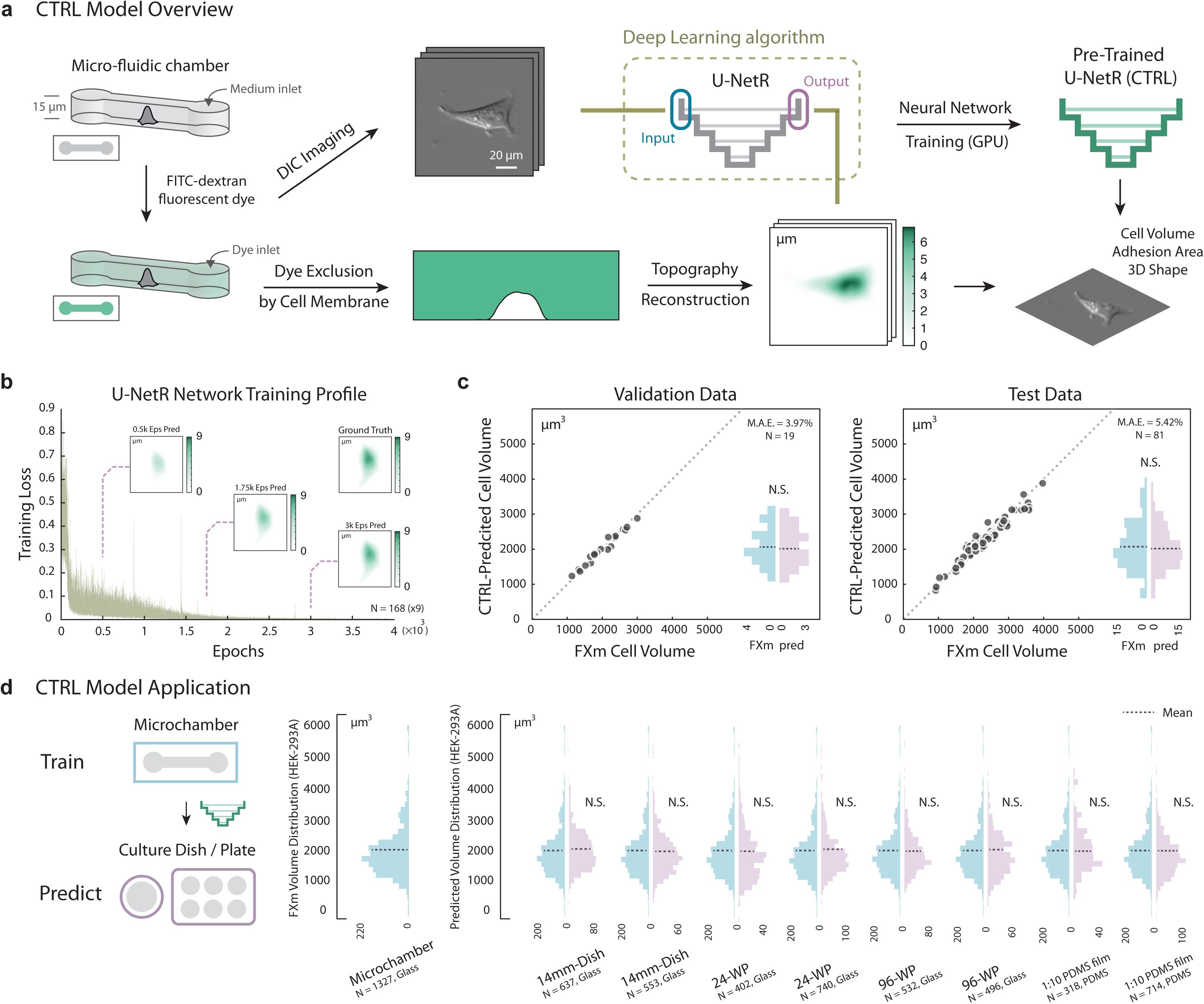
Overview of CTRL model construction, validation, and application. **(a) CTRL model overview**. Cells are seeded into a micro-fabricated chamber for the Fluorescence Exclusion method (FXm). A membrane-impermeable fluorescent dye (dextran) is injected into the chamber for dye exclusion measurement. A DIC microscopy image is acquired. A cell topography image is reconstructed from the fluorescent image simultaneously acquired for the same cell. Two such images form an image pair and serve as the input and the output in the Convolutional Neural Network: U-NetR (U-Net Regression Network). Training is performed with GPU to obtain a trained U-NetR based CTRL model. With the trained CTRL model, cell volume and topology can be computed and compared to training data as validation or test. **(b) U-NetR model training profile**. Training profile of U-NetR based on training data of 1512 images (9 augmentations for each cell) on HEK-293A cells is shown with training loss progression over 80,000 training epochs. Predicted cell topography progression over the training course is shown for a representative cell at 0.5k, 1.75k, and 3k epochs. Measured cell topography image (FXm data) for the cell is shown in the top-right corner. **(c) CTRL model (HEK-293A) validation**. Left: The trained CTRL model was applied on validation data (same microchamber, the other 20% of the cells); Right: The trained CTRL model was applied on test data (cells in a different microchamber). CTRL-predicted volume is compared to FXm measured cell volume for every single cell. Volume distributions are shown as histograms. No statistical significance is found between measured volume and predicted volume. Cell number and Mean Absolute Error (M.A.E.) for both the validation and test case are indicated in the top-right corner of each panel. **(d) CTRL model application across different cell culture platforms**. A CTRL model trained from microchamber data can be applied to DIC cell images acquired on a variety of glass and PDMS substrates, eliminating the need for repeated FXm experiment. The only input for the CTRL model is a DIC image. HEK-293A cells were seeded in a 14mm-dish, a 24-well plate, in a 96-well plate, and a 1:10 PDMS membrane (2 biological repeats, each biological repeat contains 3 technical repeats). Single-cell DIC images were inputs for the trained CTRL model. Distributions of the predicted volume in each condition are plotted in histograms with the mean indicated by a dashed line. No statistical difference is found between the CTRL-predicted cell volume distribution and the measured volume distribution in any of the test conditions.

Convolutional Neural Networks (CNN) have been applied to microscopy images for both phenotype classification^21,22^ and image segmentation^23,24^. CNNs trained on these tasks have proven to be accurate and predictive, and have demonstrated significant potential to generalize to a wide range of predictive tasks in biology. Here we present CTRL: Cell Topography Reconstruction Learner, a novel label-free technique that uses the U-Net Regression Network and the FXm for reconstructing cell topography and estimating single-cell volume from Differential Interference Contrast (DIC) microscopy images. The method requires a one-time training data set of FXm single-cell images and their corresponding DIC images. Once trained, the method can be used to predict single-cell volume to quantitative accuracy without microchambers and fluorescence labeling. The method allows for continuous single-cell volume measurements in multiple types of cell culture platforms without time limit.

The image translation CNN (UNet) was first proposed by Olaf et al.^23^. As Fig. S1 shows, the CNN takes in a digital image as the input and generates a corresponding digital image as the output. The progressive structure between the first and last layer is composed of not only traditional convolution, ReLU, and down-sampling layers, but also up-convolution and up-sampling layers, enabling the Network to “recover” a parallel output image for the given the input image (image-to-image translation). The application of U-Net in computational biology^25^ has mainly focused on image segmentation by performing pixel classification^24^. Here we modify the last layer in traditional U-Net structure and present a U-Net Network for pixel regression: U-Net Regression Network (U-NetR) (Fig. S1). As opposed to predicting a categorical label, U-NetR predicts a positive real number for each pixel. For data collection, we acquire DIC image data and the corresponding microchamber fluorescence image data from the FXm experiment (Fig. 1a). Images are taken immediately after the introduction of fluorescent dye, therefore the data excludes potential effects of endocytosis. Pre-processing of DIC images and microchamber fluorescence images and the intensity augmentation for DIC images (Fig. S2, S3) are detailed in Methods. For training data, the DIC image serves as the input and the fluorescence image serves as the output. U-NetR aims to uncover the hidden relationship between the image pair during Network training. In other words, U-NetR learns to generate the cell height map (topography) from the DIC image, given hundreds to thousands of “image pairs”. The methodology is built upon the hypothesis that the intensity distribution over a DIC image contains information in the cell height map that is inexplicable to human eyes and unsolvable via traditional image processing algorithms. Previous attempts have been made to model the optics of the DIC microscope to recover the object 3D shape^26^. Here we use the U-Net to optimize this mapping without knowledge of microscopy details.

We applied U-NetR training on HEK-293A cells (Fig. 1b), which showed a gradual decrease of training loss over 80,000 epochs. The validation and test results (Fig. 1c, Fig. S4) show that the prediction Mean Absolute Error (M.A.E.) remains less than 4% for validation data (same microchamber, the remaining 20% of the cells) and 6% for test data with 3 biological repeats (cells from a different microchamber in a separate experiment) and the absolute error of the population average is less than 3% for both cases. These results suggest that a trained CTRL model can adequately predict cell volume for unseen DIC image data. The predicted cell volume distributions are also quantitatively similar to the measured cell volume distribution from the FXm (Fig. 1c). The representative convolutional feature maps from the trained model are shown in Fig. S5. In particular, some of the feature maps show diffraction-grating-like grids (Fig. S5a), suggesting that U-NetR is learning optics. Comparing to the traditional FXm method, this computational approach achieves significant savings in laboratory work, requiring no micro-device fabrication and fluorescence imaging beyond the training data set. More importantly, it liberates cells from the confinement of microchambers and long-term incubation together with fluorescence dyes, and therefore ensures normal cell growth and eliminates errors from dye endocytosis. To test the applicability and robustness of the method to different substrates used during DIC imaging, we tested varying cell culture seeding platforms from glass to polydimethylsiloxane (PDMS) substrates and report the predicted cell volume distribution (Fig. 1d). Even though the training data was taking from microchambers with glass substrates, no significant variation is found between the predicted volume distributions of the populations across different platforms.

To further explore the applicability of the method, we investigated the effect of pharmacological inhibition on model generalization. Rapamycin, an mTOR pathway inhibitor, has been shown to decrease cell volume by ∼15%^4,20,27,28^. We measured the cell volume data of rapamycin-treated (72hrs, 1nM) HEK-293A cells from the FXm method and applied the CTRL model trained on untreated cells to the corresponding DIC images. Results show that the predicted cell volume distribution matches cell volume distribution from FXm experiments (Fig. 2a). Next we tested HEK-293A CRISPR knockout of the Hippo pathway protein YAP^29,30^, a nuclear transcription factor that regulates cell volume^31^ in an mTOR independent manner. The CTRL model (trained on WT cells) predictions are again in excellent agreement with FXm (Fig. 2b). It is also known that biomechanical environment of the cell such as the substrate stiffness affects cell volume^19,32^, and we found that the model trained on data from glass substrates (GPa) can predict cell volume on 3kPa PDMS substrates (Fig. 2d).

**Fig. 2.**
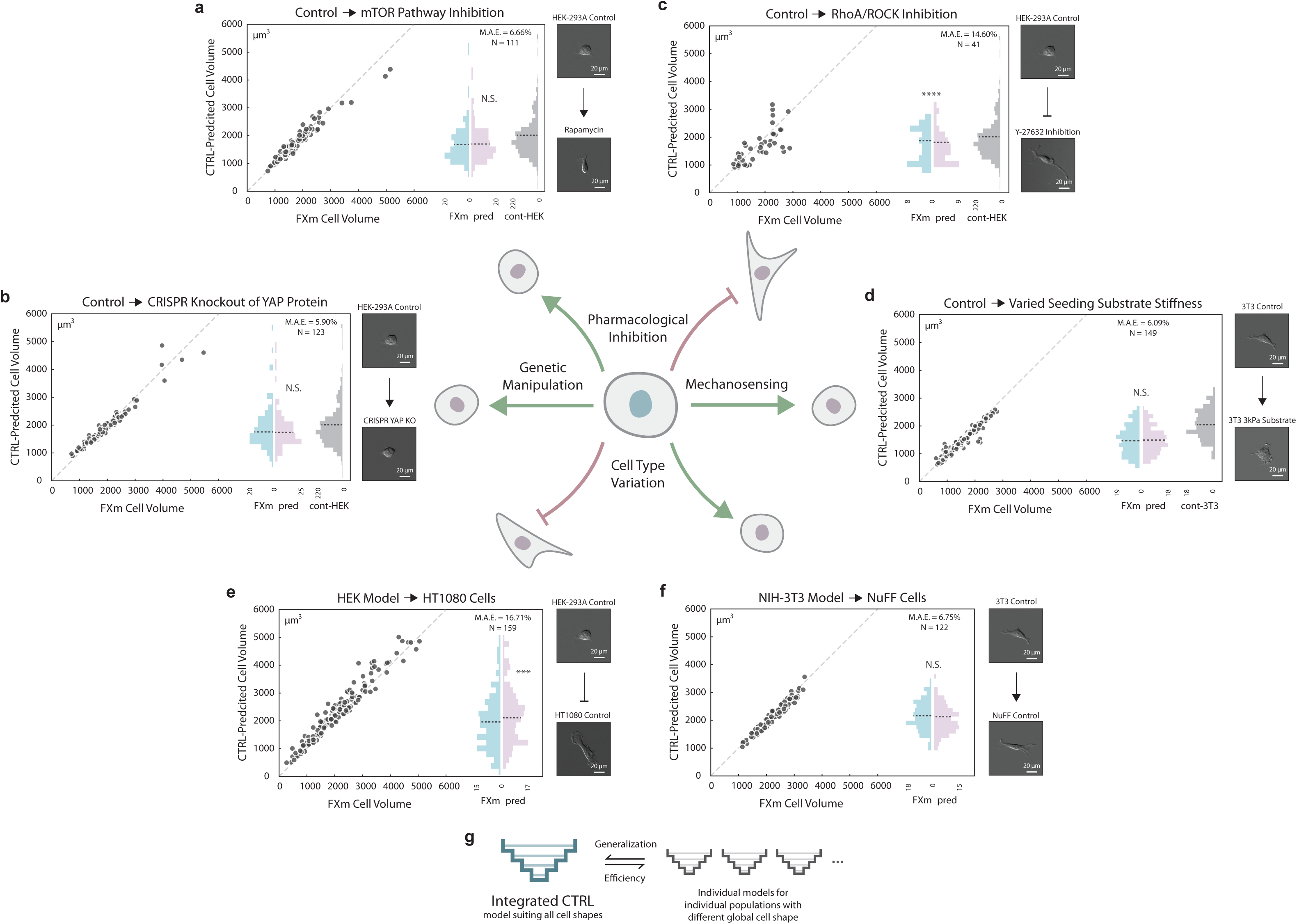
CTRL model generalization. **(a) Generalization to cells with mTOR pathway inhibition via rapamycin**. HEK-293A cells were treated with 1nM rapamycin for 72 h before the FXm experiment. Cell volume was experimentally measured via the FXm and predicted by a CTRL model previously trained on data from untreated HEK-293A cells (Fig. 1c). No statistical difference is found between the measured volume and the predicted volume distributions. Cell volume distribution of control HEK-293A cells (training data) is plotted in gray as a reference. **(b) Generalization to CRISPR knockout of YAP protein**. YAP KO of HEK-293A was generated previously^29,30^. Cell volume was measured via the FXm and predicted by a CTRL model previously trained on data from WT HEK-293A cells (Fig. 1c). No statistical difference is found between measured volume and predicted volume. Measured cell volume distribution of control HEK-293A cells (training data) is shown in gray as a reference. **(c) Generalization to cells with ROCK inhibition via Y-27632 cannot be achieved**. HEK-293A cells were treated with 100μM Y-27632 for 2 h before the FXm experiment. Cell volume was experimentally measured via the FXm and predicted by a CTRL model previously trained on data from untreated HEK-293A cells (Fig. 1c). Measured cell volume distribution of control HEK-293A cells (training data) is plotted in gray as a reference. **(d) Generalization to PDMS substrate with different stiffness**. 3T3 cells were seeded on a PDMS substrate with stiffness of 3kPa. Cell volume was experimentally measured via the FXm and predicted by a CTRL model previously trained on data from HEK-293A cells on regular glass substrates (Fig. 1c). No statistical difference is found between FXm-measured volume and predicted volume. Measured cell volume distribution of 3T3 cells on regular glass substrate (training data) is plotted in gray as a reference. **(e) Poor generalization to HT1080 cells with CTRL model trained on HEK-293A cells**. U-NetR was trained on HEK-293A cells and tested on HT1080 cells. CTRL-predicted volume is compared to FXm-measured cell volume for every single cell. **(f) Generalization to NuFF cells with CTRL model trained on NIH-3T3 cells**. U-NetR was trained on NIH-3T3 cells and tested on NuFF cells. CTRL-predicted volume is compared to FXm-measured cell volume for every single cell. **(g) Illustration of the relationship between an integrated model and individual models**. Individual CTRL models train on specific cell types may not generalize to other cell types with substantially different cell shapes, while an integrated model pooling training data from all cell types is able to achieve generalization (see Fig. S6). The trade-off for the integrated model will be greater training time. *For (a)∼(f), DIC images of a representative cell of each training population and each test population are displayed.

We then sought to see if the model generalization persisted in situations where there is a dramatic change in cell shape. Y-27632 is known to inhibit^33,34^ Rho kinase activity and cell contractility, and change the cell shape. We found that the model (trained on untreated cells) no longer can predict the volume during Y-27632 treatment (Fig. 2c) with M.A.E greater than 14%. We then asked if a CTRL model trained on one cell type can generalize to a different cell type with a different cell shape. It was found that a CTRL model trained on data acquired for HEK-293A is able to predict larger average volume of HT1080 (fibrosarcoma cells) (Fig. 2e), but the M.A.E is too large to be quantitative at the individual level. However, a CTRL model trained on data from NIH-3T3 cells is able to predict cell volume accurately for NuFF cells (newborn foreskin fibroblast) (Fig. 2f), which is of the same cell type (fibroblast) as 3T3 and has a similar cell shape. These results (Fig.2a, b, c, d, e, f) suggest that the CTRL model generalizes well to biological perturbation of the same cell type, but fails when there is a dramatic cell shape change. It is reasonable to conjecture that pooling images from multiple cell types with different shape variations together as the training data will generate a general model (Fig. 2g). Indeed, a CTRL model trained on a combined data of HEK-293A, HT1080 and NIH-3T3 cells can predict MDA-MB-231 cell volume accurately (Fig. S6). However, larger training data set also increases training time (Fig. S7) and a balance must be found between generalizability and computational time. Depending on the U-NetR structure, hyperparameter selection, the data augmentation scheme, the volume of the training data and the GPU used, the typical training time for a CTRL model can vary from days to weeks.

The proposed method is label-free (no fluorescent dye is needed) and liberates the cell from the microchamber. These advantages allow us to track single-cell volume in standard cell culture dishes for an indefinitely period with arbitrarily time resolution. The method allows us to quantify cell volumetric growth rates over multiple generations of daughter cells. In particular, to gain greater insight into cell size regulation, we sought to correlate the added cell volume with the birth volume of the cell over several generations. Here we used HT1080 cells instead of HEK-293A cells because the daughter cells readily separate from mother cells. A CTRL model was trained for HT1080 cells and the achieved accuracy is shown in Fig. S4. The HT1080 CTRL model prediction for a 9-hr time-lapse DIC movie is compared with the FXm volume measurement for the same cell (Fig. 3b). The model again demonstrates quantitative cell volume prediction even through cell division, including on the phenomena of mitotic swelling^3^. The prediction error over time (frames) for the entire population is shown in Fig. S8, showing that the CTRL model is quantitatively accurate for all cells tracked. We then applied the CTRL model for a 50hr DIC movie of growing and dividing cells in a standard cell culture dish (Fig. 3a). The cell volume trajectory of a single-cell and one of the daughter cells from each of its three divisions (generations) are shown in Fig. 3a. In order to quantify cell volume before and after cell division, it is essential to include images of cells undergoing division in the training data. Note that due to dye endocytosis, the experiment in Fig. 3a is not possible with FXm experiments. Results of added cell volume during a cell cycle versus the cell volume at birth (Fig. 3c) show that HT1080 adopts a sizer-like growth mechanism. The time-lapse experiment also revealed the correlation between the cell cycle length (division time), volume growth rate and cell birth volume (Fig. S9). Moreover, the coefficient of variation, 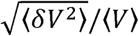, is generally constant over the cell cycle, but shows a visible decrease at 25% cell cycle completion (Fig. 3e). This is indicative of a cell cycle checkpoint where size-control would reduce cell size fluctuations (Fig. 3d). The CTRL model is also useful for quantifying rapid cell volume changes such as during an osmotic shock (Fig. 3f). Here the cell volume is tracked every 30s, showing HT1080 cells can re-adjust their volumes after a hypotonic shock (50%) in 15-30min (single-cell trajectories are shown in Fig. S10).

**Fig. 3.**
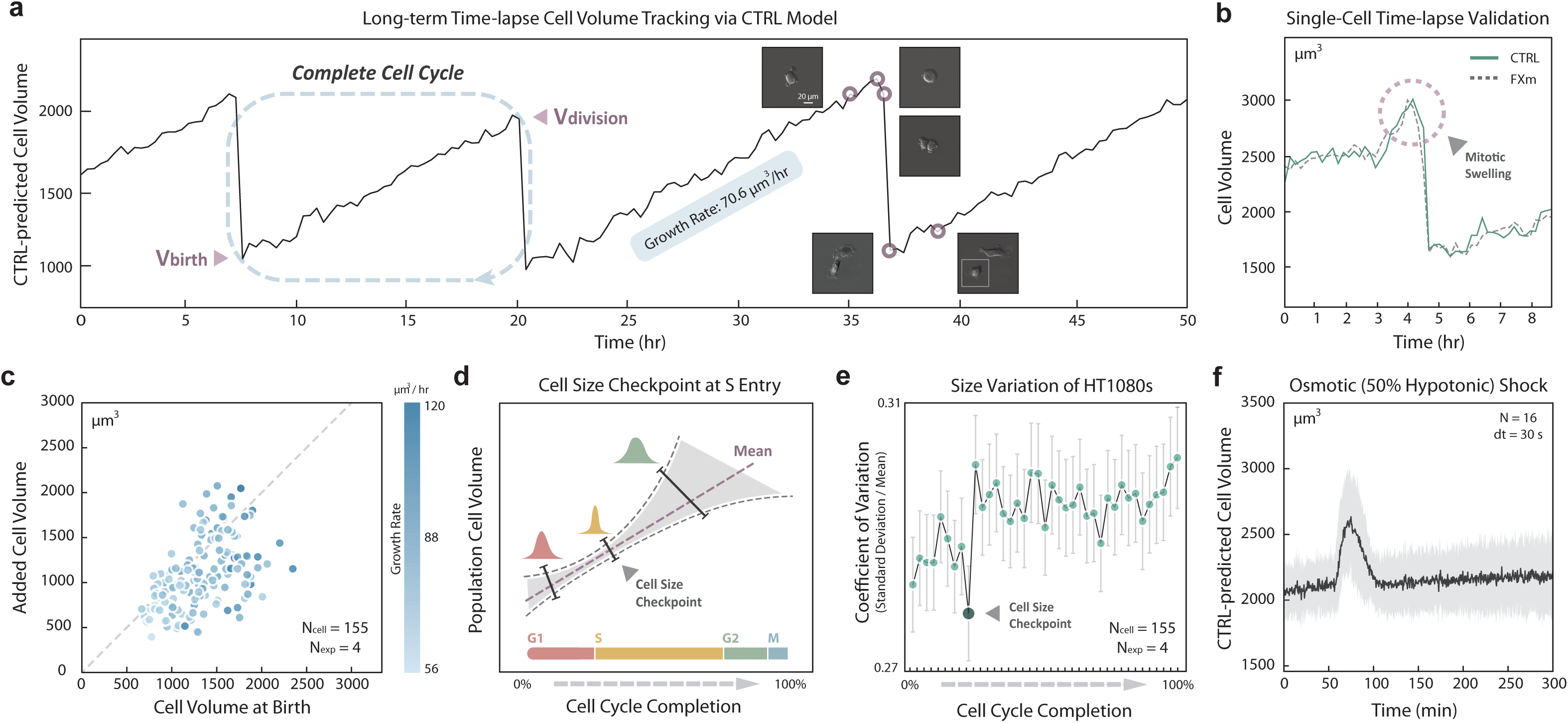
Time-lapse cell volume tracking via CTRL. **(a) Representative long-term cell volume trajectory**. Cell volume from CTRL prediction of a single HT1080 cell is plotted over time (50 h) with 3 visible divisions. DIC images of the cell before and after division at several time points are shown. The growth rate within one cell cycle is displayed. A linear growth law is assumed. **(b) Time-lapse single-cell volume validation**. Cell volume of HT1080 cells was quantified using CTRL model (solid) on DIC images and compared to experimental data from the FXm (dashed line) for the same cell. The time interval between adjacent frames is 20 min. **(c) The relationship between added cell volume and cell volume at birth**. From predicted HT1080 time-lapse cell volume trajectories, the added cell volume for one cell cycle and is plotted against the cell volume at birth. To ensure that each cell has completed a full cell cycle, cells with two consecutive visible divisions are analyzed. The dotted line indicates added cell volume equal to birth volume. Data is from 4 biological repeats (N=155). The data indicate that HT1080 is a sizer. **(d) Cell size checkpoint**. The presence of a cell size checkpoint is a potential mechanism for maintaining size homeostasis for the entire population. Cells progress to a new cell cycle phase, e.g. S entry, only when the physical size of the cell reaches a size threshold (checkpoint). If there is such a checkpoint, cell size variation at the checkpoint should decrease due to feedback control. **(e) Cell volume coefficient of variation (CV) for the complete cell cycle**. Individual single-cell volume was tracked for >70h. To ensure each cell has completed a full cell cycle, only cells with two consecutive visible divisions are analyzed. The cell cycle is divided into 39 increments (2.5% for each increment), and we analyzed the collected volumes of all cells at each increment of cell cycle completion. The mean (Fig. S9c) and the standard deviation (Fig. S9d) of the cell volume, and the coefficient of variation (CV = standard deviation / mean) are shown. A visible decrease of CV was found at 25% cell cycle completion, indicating the presence of cell size checkpoint for HT1080s. Error bars represent standard deviations of CV generated from 1000 random sampling of 60 cells from available 155 cells. **(f) Cell volume adaptation during osmotic shock**. 50% hypotonic shock medium was added to HT1080 cells. DIC images were taken 60 min before the shock and 240 min after the shock with high time resolution (30 s). Averages cell volume over all single cells in 2 biological repeats (N=19) is shown in black line, and the standard deviation over all timepoints is supplied (gray interval). Single-cell volume trajectories are displayed in Fig. S10.

Understanding the mechanisms controlling cell size and cell growth is a fundamental goal in cell and tissue biology. So far, the lack of a label-free and easy-to-use techniques that can accurately measure cell volume has limited quantitative studies on factors influencing mammalian cell size and cell growth dynamics. The present methodology is quantitatively accurate in estimating both static and time-lapse single-cell size in standard cell culture conditions in a high throughput manner. The method requires a one-time data collection of training images using the FXm method, but subsequent volume predictions can be done in standard cell culture dishes without labeling. The method can also be generalized to any cell type with enough input training data, perhaps from a global database. More generally, Artificial Intelligence methods can provide new insights and opportunities for image-based biological discovery while reducing experimental time and cost.

## Acknowledgments

This work has been funded in part by National Institutes of Health grants U54CA210172 and R01GM134542.

## Author contributions

K.Y., N. D. R., and S. X. S designed the study. K. Y. performed all experiments, Convolutional Neural Network design and training, and data analysis. K.Y. and S. X. S. wrote the paper. S.X.S. supervised the study.

## Data availability

All original data in the present work is available from the corresponding author upon reasonable request.

## Code availability

All MATLAB codes written and applied in Convolutional Neural Network design and training, image pre-processing and validation analyses, as well as exemplary image data, are available online at https://GitHub.com/sxslabjhu/.

## Competing interests

The authors declare no conflict of interest.

## Notes

https://github.com/sxslabjhu/CTRL/

## References

1. Ginzberg, M. B., Kafri, R. & Kirschner, M. On being the right (cell) size. Science 348, (2015).

2. Lloyd, A. C. The regulation of cell size. Cell 154, 1194 (2013).

3. Zlotek-Zlotkiewicz, E., Monnier, S., Cappello, G., Le Berre, M. & Piel, M. Optical volume and mass measurements show that mammalian cells swell during mitosis. J. Cell Biol. (2015). doi:10.1083/jcb.201505056

4. Pollizzi, K. N., Waickman, A. T., Patel, C. H., Sun, I. H. & Powell, J. D. Cellular size as a means of tracking mTOR activity and cell fate of CD4+ T cells upon antigen recognition. PLoS One (2015). doi:10.1371/journal.pone.0121710

5. Björklund, M. Cell size homeostasis: Metabolic control of growth and cell division. Biochimica et Biophysica Acta - Molecular Cell Research (2019). doi:10.1016/j.bbamcr.2018.10.002

6. Edens, L. J., White, K. H., Jevtic, P., Li, X. & Levy, D. L. Nuclear size regulation: From single cells to development and disease. Trends in Cell Biology (2013). doi:10.1016/j.tcb.2012.11.004

7. Stenkula, K. G. & Erlanson-Albertsson, C. Adipose cell size: importance in health and disease. Am. J. Physiol. Regul. Integr. Comp. Physiol. 315, R284–R295 (2018).

8. Kozma, S. C. & Thomas, G. Regulation of cell size in growth, development and human disease: PI3K, PKB and S6K - Kozma - 2002 - BioEssays - Wiley Online Library. Bioessays (2002). doi:10.1002/bies.10031

9. Si, F. et al. Mechanistic Origin of Cell-Size Control and Homeostasis in Bacteria. Curr. Biol. 29, 1760-1770.e7 (2019).

10. Hevia, D. et al. Cell volume and geometric parameters determination in living cells using confocal microscopy and 3D reconstruction. Protoc. Exch. (2011). doi:10.1038/protex.2011.272

11. Du, T. & Wasser, M. 3D Image stack reconstruction in liwe cell microscopf of drosophila muscles and its validation. Cytom. Part A (2009). doi:10.1002/cyto.a.20701

12. Hirsch, J. & Gallian, E. Methods for the determination of adipose cell size in man and animals. J. Lipid Res. (1968).

13. Kubitschek, H. E. & Friske, J. A. Determination of bacterial cell volume with the Coulter Counter. J. Bacteriol. (1986). doi:10.1128/jb.168.3.1466-1467.1986

14. Gray, M. L., Hoffman, R. A. & Hansen, W. P. A new method for cell volume measurement based on volume exclusion of a fluorescent dye. Cytometry (1983). doi:10.1002/cyto.990030607

15. Stern, A. D., Rahman, A. H. & Birtwistle, M. R. Cell size assays for mass cytometry. Cytom. Part A (2017). doi:10.1002/cyto.a.23000

16. Tzur, A., Moore, J. K., Jorgensen, P., Shapiro, H. M. & Kirschner, M. W. Optimizing optical flow cytometry for cell volume-based sorting and analysis. PLoS One (2011). doi:10.1371/journal.pone.0016053

17. Cermak, N. et al. High-throughput measurement of single-cell growth rates using serial microfluidic mass sensor arrays. Nat. Biotechnol. 34, 1052–1059 (2016).

18. Cadart, C. et al. Fluorescence eXclusion Measurement of volume in live cells. Methods Cell Biol. 139, 103–120 (2017).

19. Gonzalez, N. P. et al. Cell tension and mechanical regulation of cell volume. Mol. Biol. Cell (2018). doi:10.1091/mbc.E18-04-0213

20. Perez-Gonzalez, N. A. et al. YAP and TAZ regulate cell volume. J. Cell Biol. jcb.201902067 (2019). doi:10.1083/jcb.201902067

21. Kihm, A., Kaestner, L., Wagner, C. & Quint, S. Classification of red blood cell shapes in flow using outlier tolerant machine learning. PLoS Comput. Biol. (2018). doi:10.1371/journal.pcbi.1006278

22. Yao, K., Rochman, N. D. & Sun, S. X. Cell Type Classification and Unsupervised Morphological Phenotyping From Low-Resolution Images Using Deep Learning. Sci. Rep. 9, 13467 (2019).

23. Ronneberger, O., Fischer, P. & Brox, T. U-net: Convolutional networks for biomedical image segmentation. in Lecture Notes in Computer Science (including subseries Lecture Notes in Artificial Intelligence and Lecture Notes in Bioinformatics) (2015). doi:10.1007/978-3-319-24574-4_28

24. Ibtehaz, N. & Rahman, M. S. MultiResUNet□: Rethinking the U-Net Architecture for Multimodal Biomedical Image Segmentation. (2019).

25. Falk, T. et al. U-Net: deep learning for cell counting, detection, and morphometry. Nat. Methods (2019). doi:10.1038/s41592-018-0261-2

26. Kagalwala, F. & Kanade, T. Reconstructing specimens using DIC microscope images. IEEE Trans. Syst. Man Cybern. Part B 33, 728–737 (2003).

27. Fingar, D. C., Salama, S., Tsou, C., Harlow, E. & Blenis, J. Mammalian cell size is controlled by mTOR and its downstream targets S6K1 and 4EBP1/eIF4E. Genes Dev. (2002). doi:10.1101/gad.995802

28. Inoki, K., Ouyang, H., Li, Y. & Guan, K.-L. Signaling by Target of Rapamycin Proteins in Cell Growth Control. Microbiol. Mol. Biol. Rev. (2005). doi:10.1128/mmbr.69.1.79-100.2005

29. Plouffe, S. W. et al. Characterization of Hippo Pathway Components by Gene Inactivation. Mol. Cell 64, 993–1008 (2016).

30. Plouffe, S. W. et al. The Hippo pathway effector proteins YAP and TAZ have both distinct and overlapping functions in the cell. J. Biol. Chem. (2018). doi:10.1074/jbc.RA118.002715

31. Gonzalez, N. A. P. et al. YAP/TAZ as a Novel Regulator of cell volume. bioRxiv (2019). doi:10.1101/528133

32. Guo, M. et al. Cell volume change through water efflux impacts cell stiffness and stem cell fate. Proc. Natl. Acad. Sci. U. S. A. 114, E8618–E8627 (2017).

33. Claassen, D. A., Desler, M. M. & Rizzino, A. ROCK inhibition enhances the recovery and growth of cryopreserved human embryonic stem cells and human induced pluripotent stem cells. Mol. Reprod. Dev. (2009). doi:10.1002/mrd.21021

34. Horani, A., Nath, A., Wasserman, M. G., Huang, T. & Brody, S. L. Rho-associated protein kinase inhibition enhances airway epithelial basal-cell proliferation and lentivirus transduction. Am. J. Respir. Cell Mol. Biol. (2013). doi:10.1165/rcmb.2013-0046TE

