## Supplementary figures and images for "CTRL: a label-free method for dynamic measurement of single-cell volume"

### Figure S1

#### Fig. S1

**a** U-NetR Structure

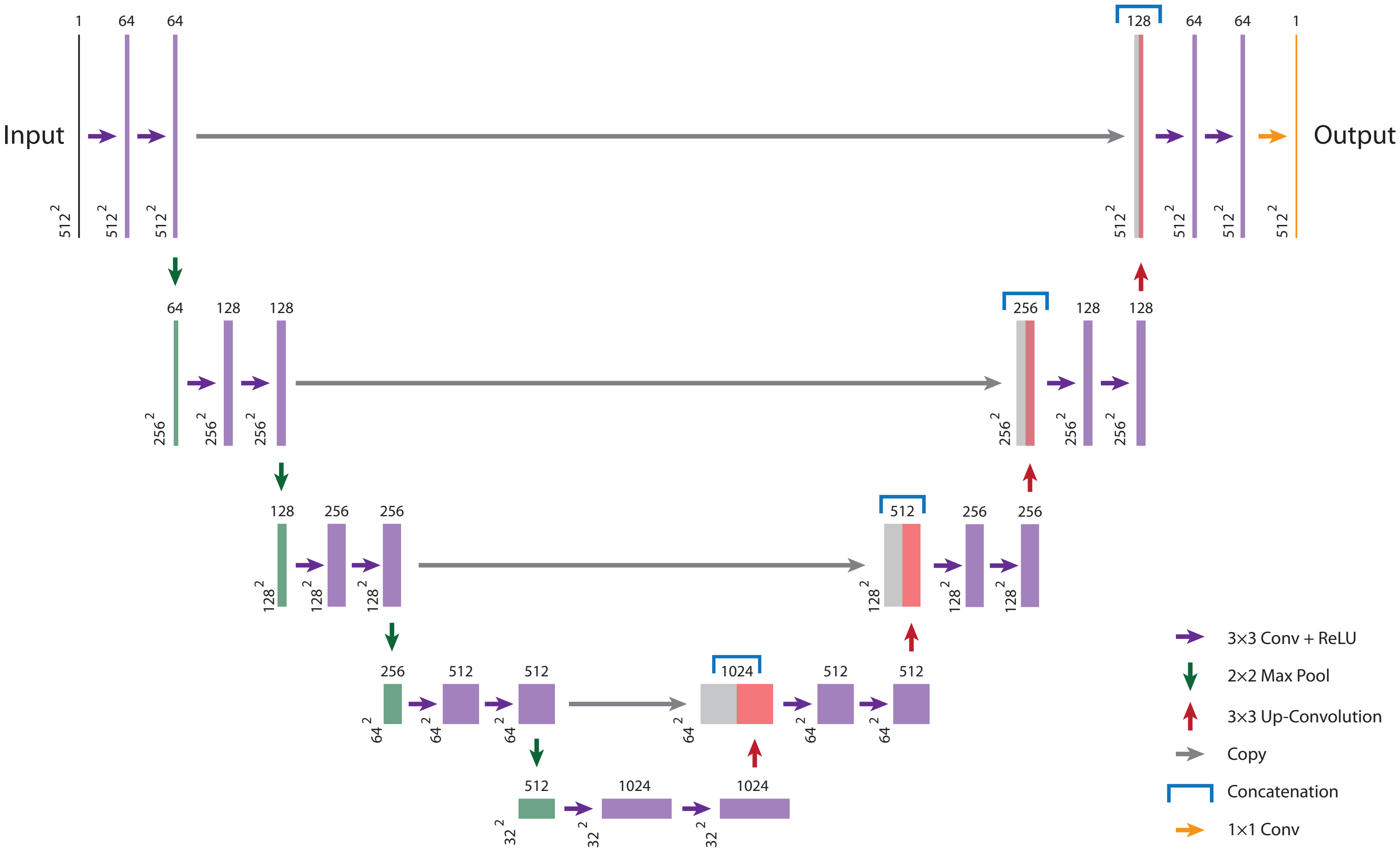

### Figure S2

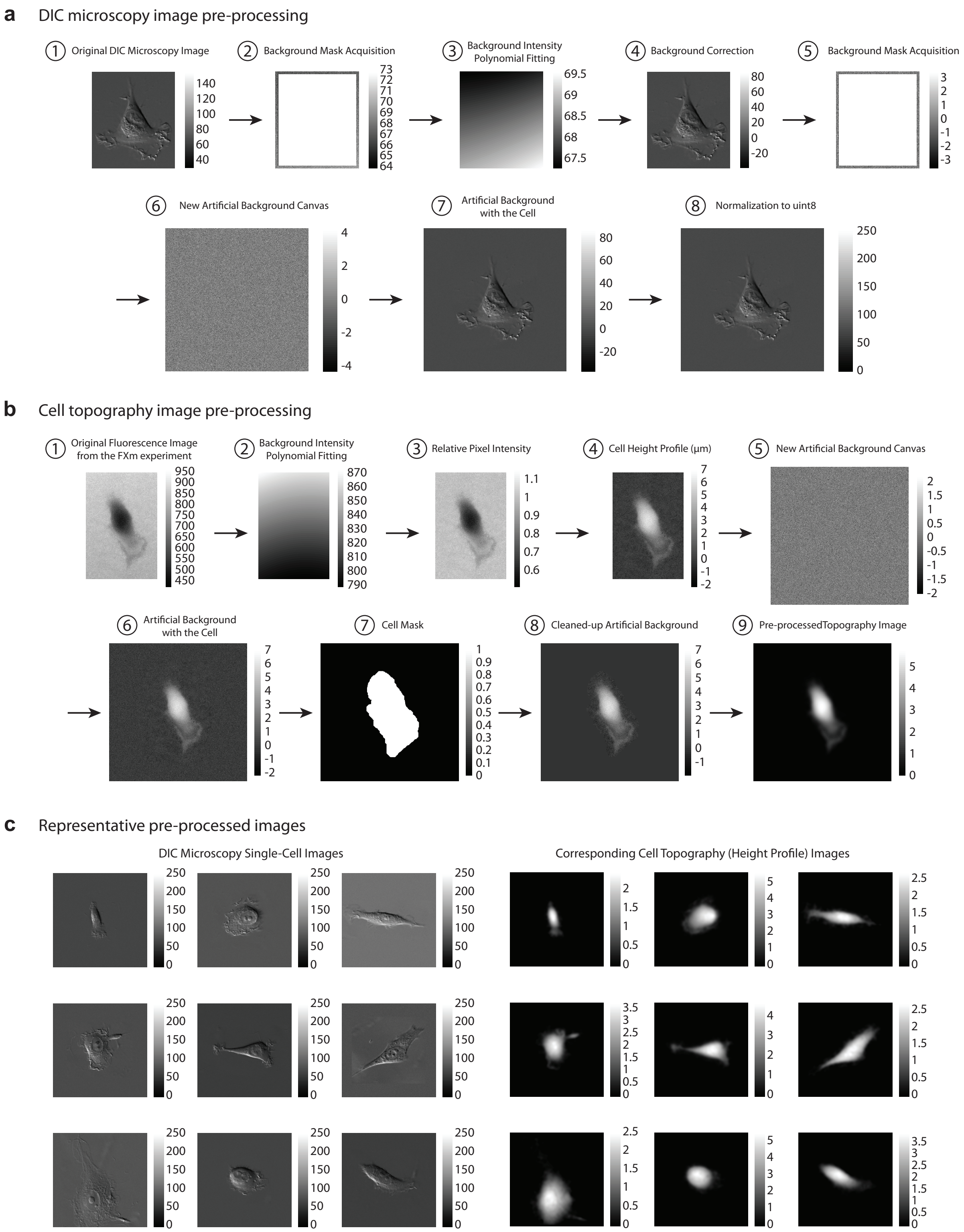

### Figure S4

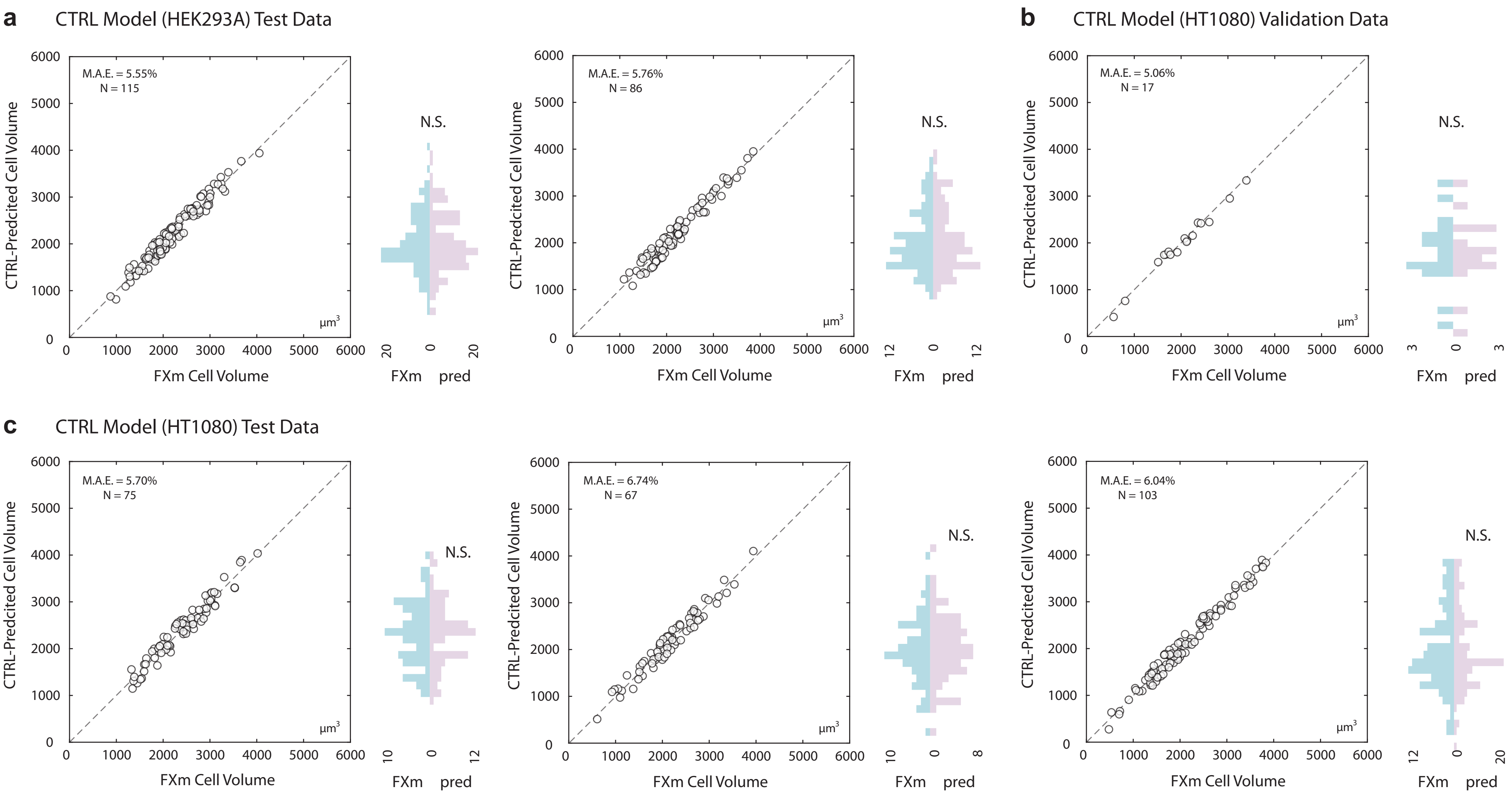

### Figure S6

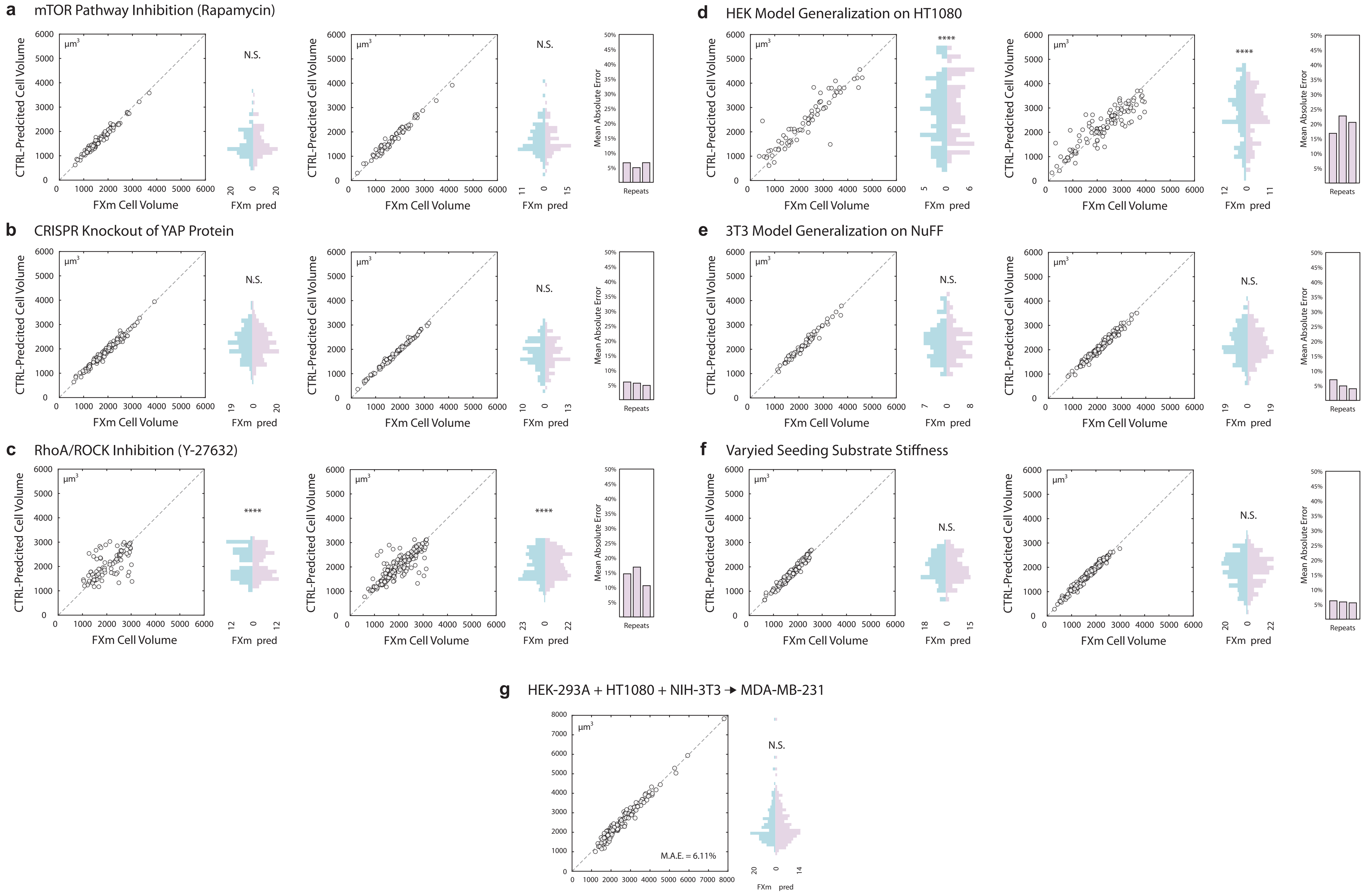

### Figure S7

Fig. S7

**a** Relationship between U-NetR training time and training data size

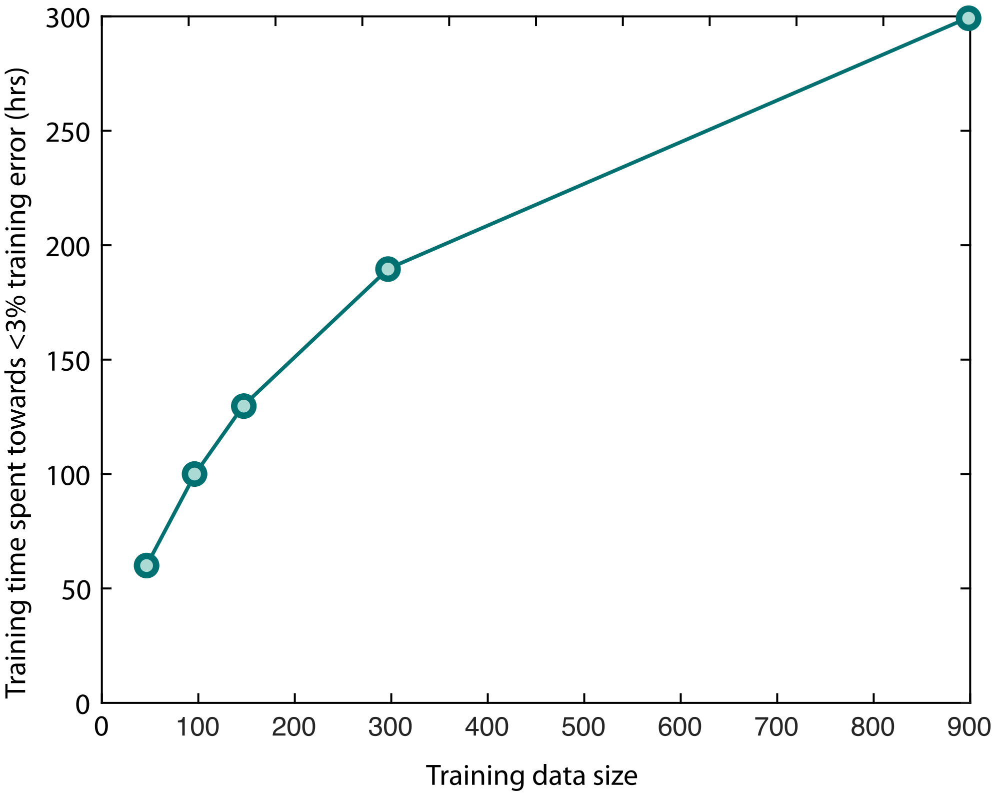

### Figure S8

Fig. S8

**a** CTRL prediction error over time in a HT1080 movie

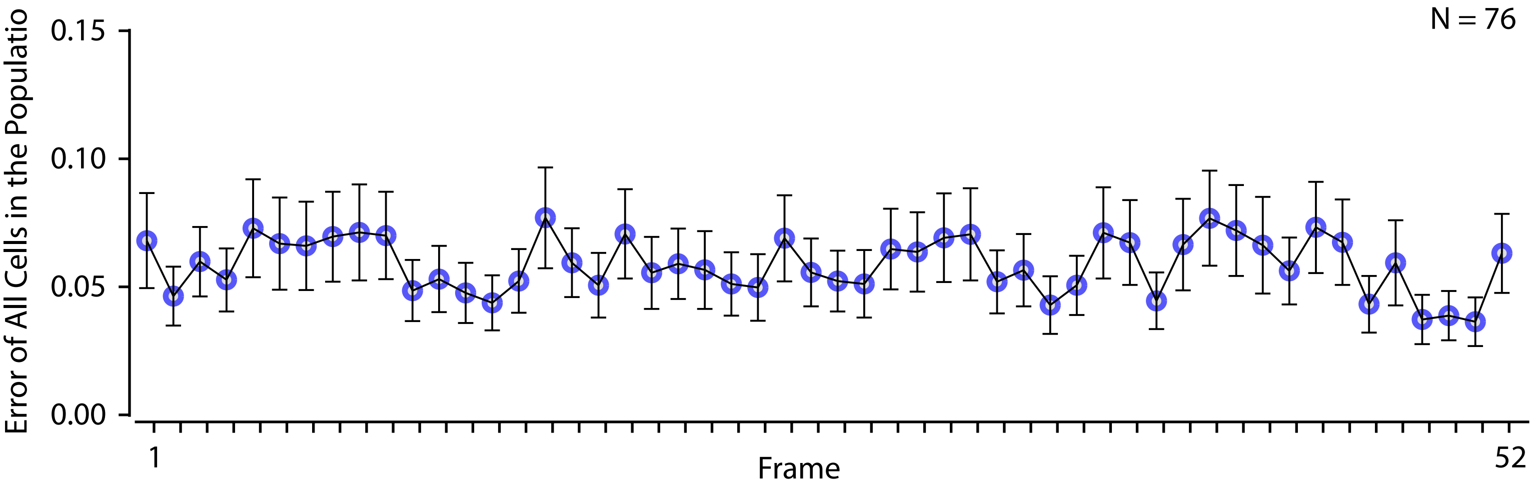

### Figure S9

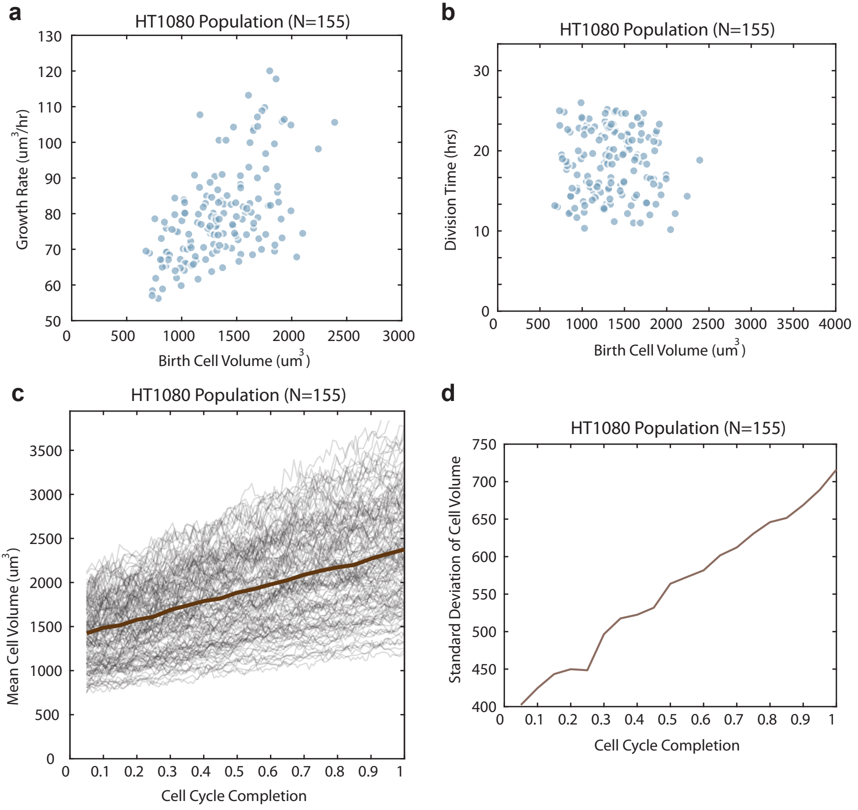

### Figure S10

#### Fig. S10

**a** Single-cell volume trajectories from osmotic shock investigation

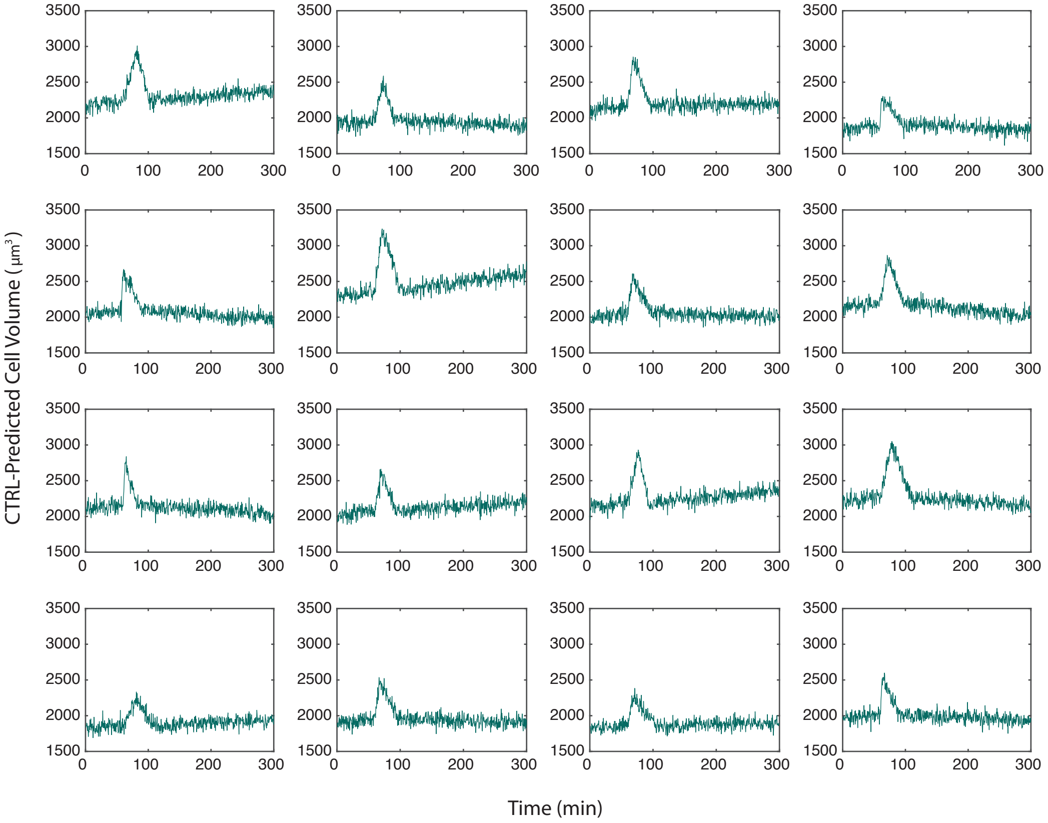
