## Supplementary material for "CTRL: a label-free method for dynamic measurement of single-cell volume": Figure S3

### Fig. S3

**a** DIC image (pre-processed) rendering for different lamp voltages

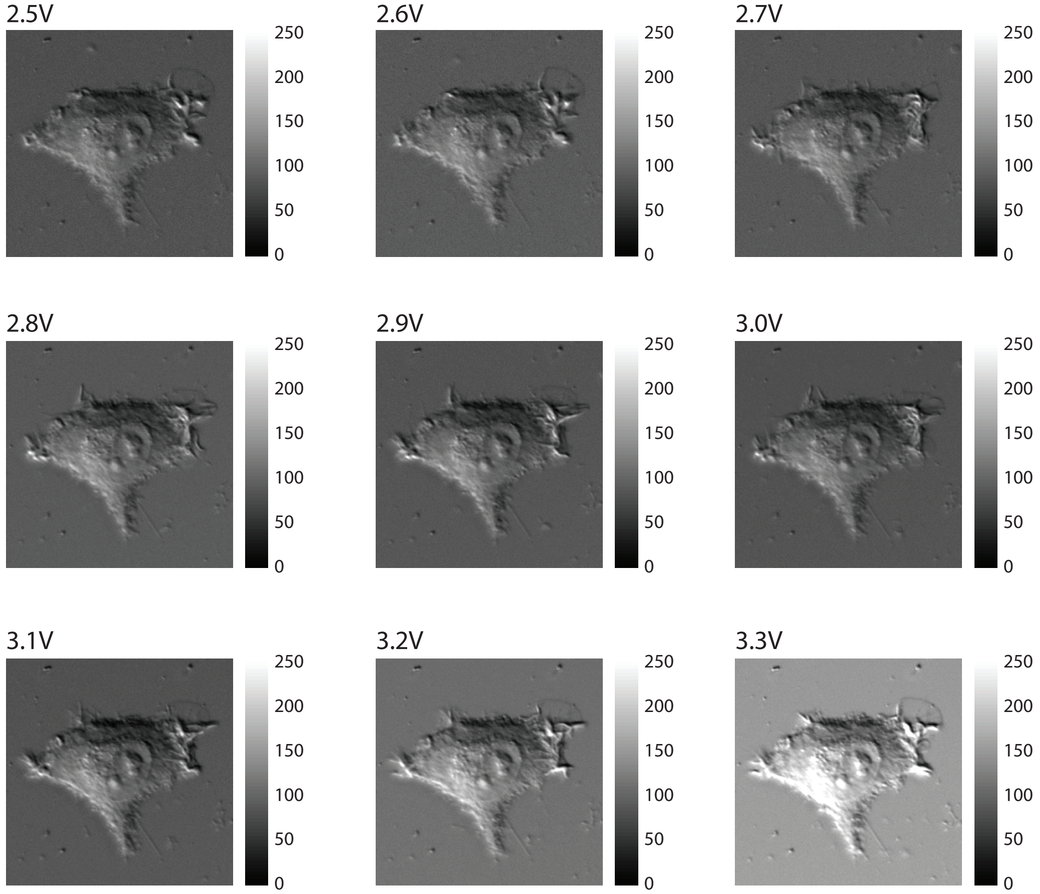


**b** DIC image histogram for different lamp voltages

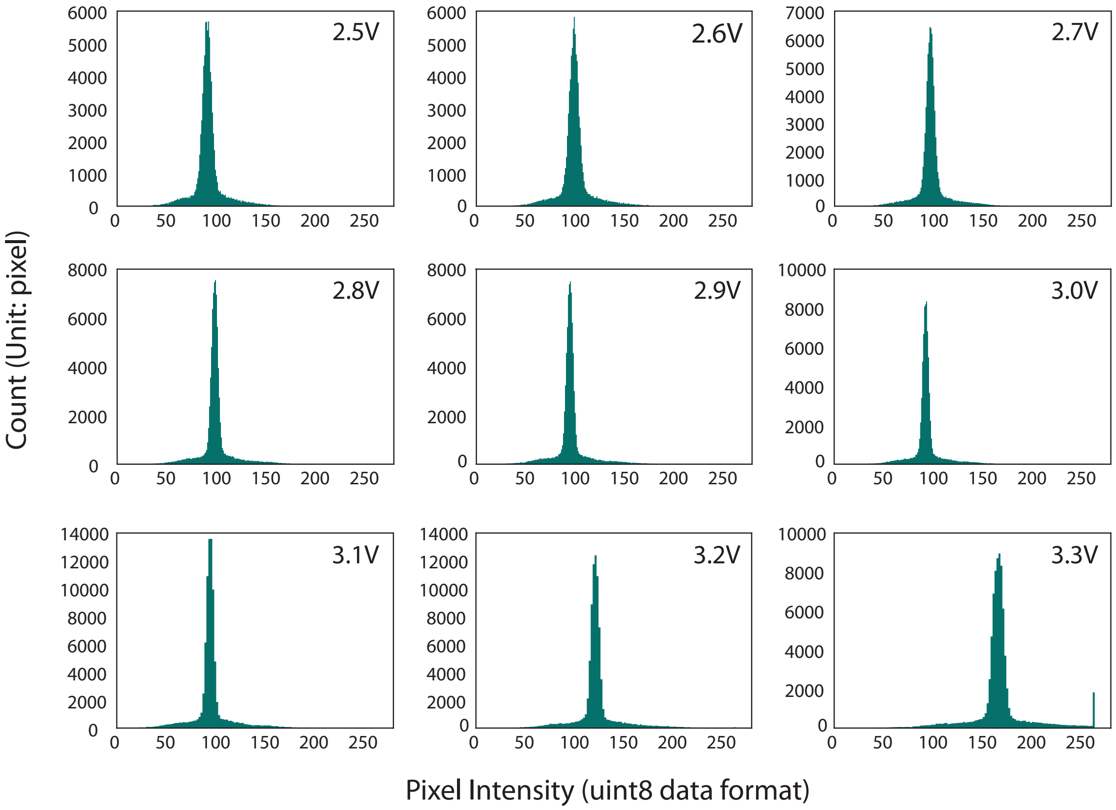
