## Supplementary material for "CTRL: a label-free method for dynamic measurement of single-cell volume": Figure S5

### Fig. S5

**a** Representative 9 feature maps (HEK-293A model, layer name: Decoding-Stage4-UpConv )

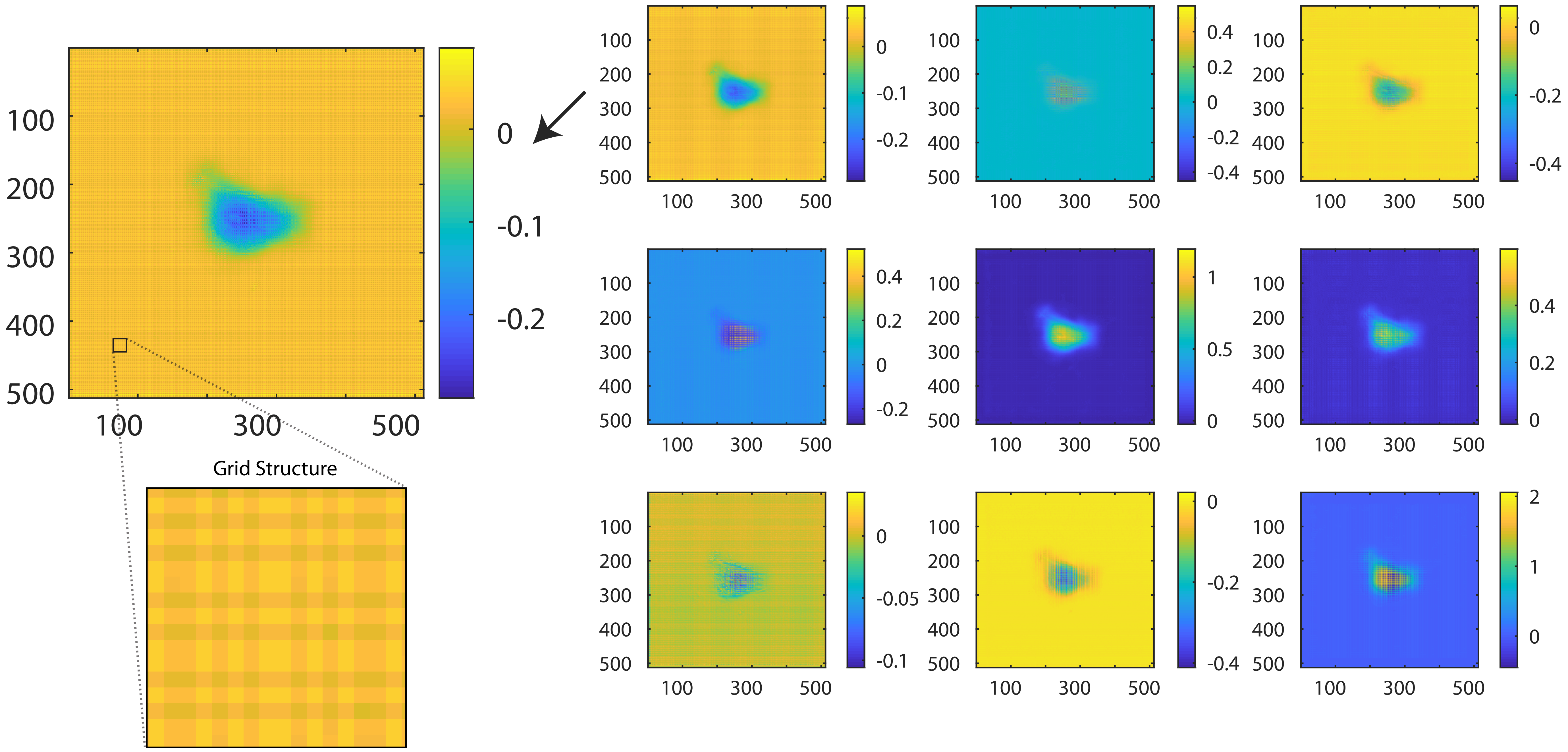


**b** Representative 9 feature maps (HEK-293A model, layer name: Decoding-Stage4-Conv1)

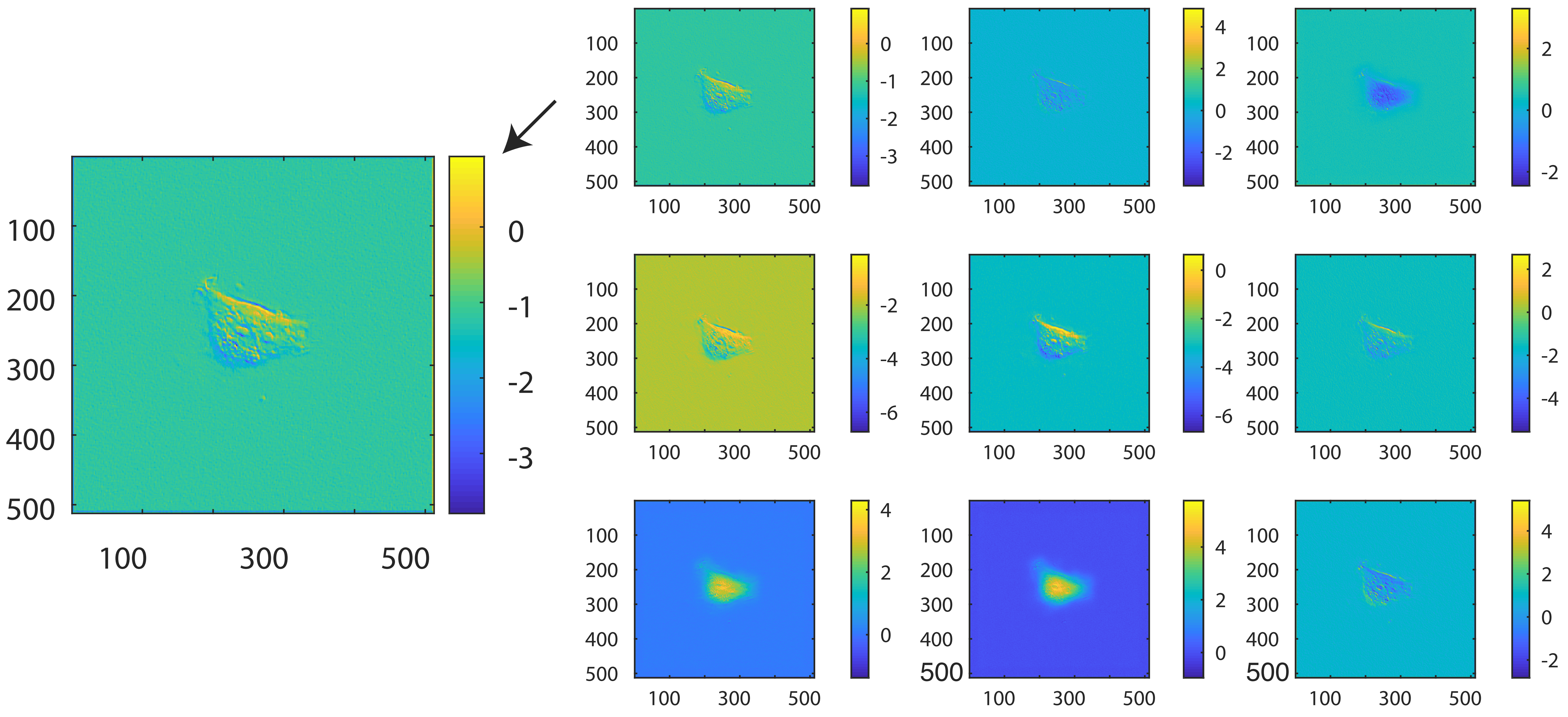
